# Evolutionary flexibility in routes to mat formation by *Pseudomonas*

**DOI:** 10.1101/2021.08.03.454897

**Authors:** Anuradha Mukherjee, Jenna Gallie

## Abstract

Many bacteria form mats at the air-liquid interface of static microcosms. These structures typically involve the secretion of exopolysaccharide(s), the production of which is often controlled by the secondary messenger c-di-GMP. Mechanisms of mat formation have been particularly well characterized in *Pseudomonas fluorescens* SBW25; mutations that lead to an increase in c-di-GMP production by diguanylate cyclases (WspR, AwsR, or MwsR) result in the secretion of cellulose, and mat formation. Here, we characterize and compare mat formation in two close relatives of SBW25: *Pseudomonas simiae* PICF7 and *Pseudomonas fluorescens* A506. We find that PICF7 – the strain more closely related to SBW25 – can form mats through mutations affecting the activity of the same three diguanylate cyclases as SBW25. However, instead of cellulose, these mutations activate the production of the Pel exopolysaccharide. We also provide evidence for at least two further – as yet uncharacterized – routes to PICF7 mat formation. *P. fluorescens* A506, while retaining the same mutational routes to mat formation as SBW25 and PICF7, forms mats by a semi-heritable mechanism that likely culminates in Pga and/or Psl production. Overall, our results demonstrate a high level of evolutionary flexibility in the molecular and structural routes to mat formation, even among close relatives.

## INTRODUCTION

Many bacteria can form structures on solid surfaces (biofilms) or at the air-liquid interface of static microcosms (mats or pellicles). The formation of these structures is dependent on the production and excretion of <u>e</u>xo<u>p</u>oly<u>s</u>accharides (EPSs) by the bacterial cells. EPS production can afford bacterial cells protection against stresses such as desiccation or starvation (Davey and O’Toole, 2000), and biofilms have been shown to increase pathogen persistence in the face of the host immune system and treatment regimens (Costerton, 1999). Given the prevalence of biofilms and mats, the evolutionary and molecular mechanisms behind their production are of interest. Here, we characterize mat production in two biocontrol agents, *Pseudomonas simiae* PICF7 and *Pseudomonas fluorescens* A506, and contrast them with previously well studied routes in *Pseudomonas fluorescens* SBW25.

The process of mat and biofilm formation often involves the concerted production of multiple structural components, and hence is under complex control. The biosynthesis of many structural components is regulated by the secondary messenger bis(3′-5′)-<u>c</u>yclic <u>di</u>meric <u>g</u>uanosine <u>m</u>ono<u>p</u>hosphate (c-di-GMP) (Ross *et al.*, 1987; Römling *et al.*, 2013; Liang, 2015). C-di-GMP is produced from GTP by enzymes called <u>d</u>i<u>g</u>uanylate <u>c</u>yclases (DGCs) with active GGDEF domains (Tal *et al.*, 1998). Conversely, c-di-GMP is broken down by <u>p</u>hospho<u>d</u>i<u>e</u>sterases (PDEs), the activity of which is dependent on conserved EAL or HD-GYP domains (Schmidt *et al.*, 2005). Most bacteria encode multiple DGCs and PDEs, the combined activities of which determine local and global intracellular c-di-GMP levels.

DGC and PDE activity is regulated by a range of stimuli, using various signal transduction pathways. A common mechanism of signal transduction in bacteria is the two-component regulatory system (Nixon *et al.*, 1986; Stock *et al.*, 2000). In such systems, a signal activates the first component (a histidine kinase), which passes the signal by a phosphotransfer event to the second component (a response regulator). Response regulators include several known examples of c-di-GMP metabolizing enzymes, including WspR of *P. fluorescens* (Bantinaki *et al.*, 2007; McDonald *et al.*, 2009) and PleD of *Caulobacter crescentus* (Paul, 2004; Chan *et al.*, 2004). In both cases, activation of a two-component signal transduction pathway results in c-di-GMP production and, ultimately, a switch from motility to a non-motile, sessile lifestyle.

The downstream targets of c-di-GMP are notably varied; known examples fall into four categories: (i) PilZ domains (Amikam and Galperin, 2006; Ryjenkov *et al.*, 2006; Lee *et al.*, 2007), (ii) degenerate, enzymatically inactive GGDEF/EAL domains (Duerig *et al.*, 2009; Colvin *et al.*, 2012; Li *et al.*, 2012), (iii) other, distinct c-di-GMP binding domains in transcriptional regulators (Hickman and Harwood, 2008; Baraquet *et al.*, 2012), and (iv) mRNA riboswitches (Sudarsan *et al.*, 2008; Chen *et al.*, 2011). Overall, c-di-GMP regulates a plethora of targets, at the levels of transcription, translation, and enzymatic activity. Many EPSs are downstream targets of c-di-GMP, including cellulosic polymers in *Escherichia coli, Acetobacter xylinum,* and *P. fluorescens* (Ross *et al.*, 1987; Goymer *et al.*, 2006; Richter *et al.*, 2020), Pga in *E. coli* and *P. fluorescens* (Boehm *et al.*, 2009; Steiner *et al.*, 2012; Lind *et al.*, 2017), and Pel in *Pseudomonas aeruginosa* and *Bacillus cereus* (Hickman and Harwood, 2008; Whitfield *et al.*, 2020).

*P. fluorescens* SBW25 is a model evolutionary system in which the mechanisms of mat formation have been particularly well characterized. SBW25 readily forms mats, or pellicles, at the air-liquid interface of static microcosms (Rainey and Travisano, 1998). The major structural component of these mats is an acetylated cellulosic polymer (Spiers *et al.*, 2003) or, when cellulose is unavailable, Pga (Lind *et al.*, 2017). Both EPSs are produced in response to high levels of c-di-GMP (Goymer *et al.*, 2006; Malone *et al.*, 2007; Lind *et al.*, 2017). In the case of cellulose, c-di-GMP is thought to target the biosynthetic machinery (Wss) by binding to the PilZ domain of WssB. The mechanism by which c-di-GMP activates SBW25 Pga has not been elucidated, but may resemble the *E. coli* mechanism of c-di-GMP binding to, and activating, Pga pathway components (Boehm *et al.*, 2009; Steiner *et al.*, 2012). In laboratory SBW25 populations, elevated c-di-GMP levels are caused by mutations affecting the activity of one of three DGCs/PDEs: WspR (DGC), AwsR (DGC), MwsR (DGC/PDE) (Bantinaki *et al.*, 2007; McDonald *et al.*, 2009; Gallie *et al.*, 2019; Lind *et al.*, 2019). The environmental signals that activate these pathways in the absence of mutation remain unknown.

In this work, we characterize and compare mat formation in two relatives of *P. fluorescens* SBW25: *P. simiae* PICF7 and *P. fluorescens* A506. We find that, despite their relatively close evolutionary relationship, each strain produces mats by distinct molecular and structural mechanisms.

## RESULTS

### Evolutionary relationships: PICF7 is more closely related to SBW25 than is A506

In this study we concentrate on three *Pseudomonas* strains: (i) *P. fluorescens* SBW25, isolated from a sugar beet leaf in the United Kingdom, (ii) *P. simiae* PICF7 (formerly *P. fluorescens* PICF7), isolated from the roots of an olive plant in Spain, and (iii) *P. fluorescens* A506, isolated from a pear plant in the USA (Table S1). All three strains have a symbiotic relationship with plants, and PICF7 and A506 are biocontrol agents for Verticillum wilt of olive (Mercado-Blanco *et al.*, 2004) and fire blight of pear and apple (Stockwell *et al.*, 2010), respectively. Complete genome sequences are available (Silby *et al.*, 2009; Loper *et al.*, 2012; Martínez-García *et al.*, 2015), and A506 also contains a sequenced, ~57 kb conjugative plasmid (pA506; Stockwell *et al.*, 2013). Phylogenetic analyses indicate that *P. fluorescens* SBW25 and *P. simiae* PICF7 more recently shared a common ancestor with each other than with *P. fluorescens* A506 (Figure 1).

**Figure 1.**
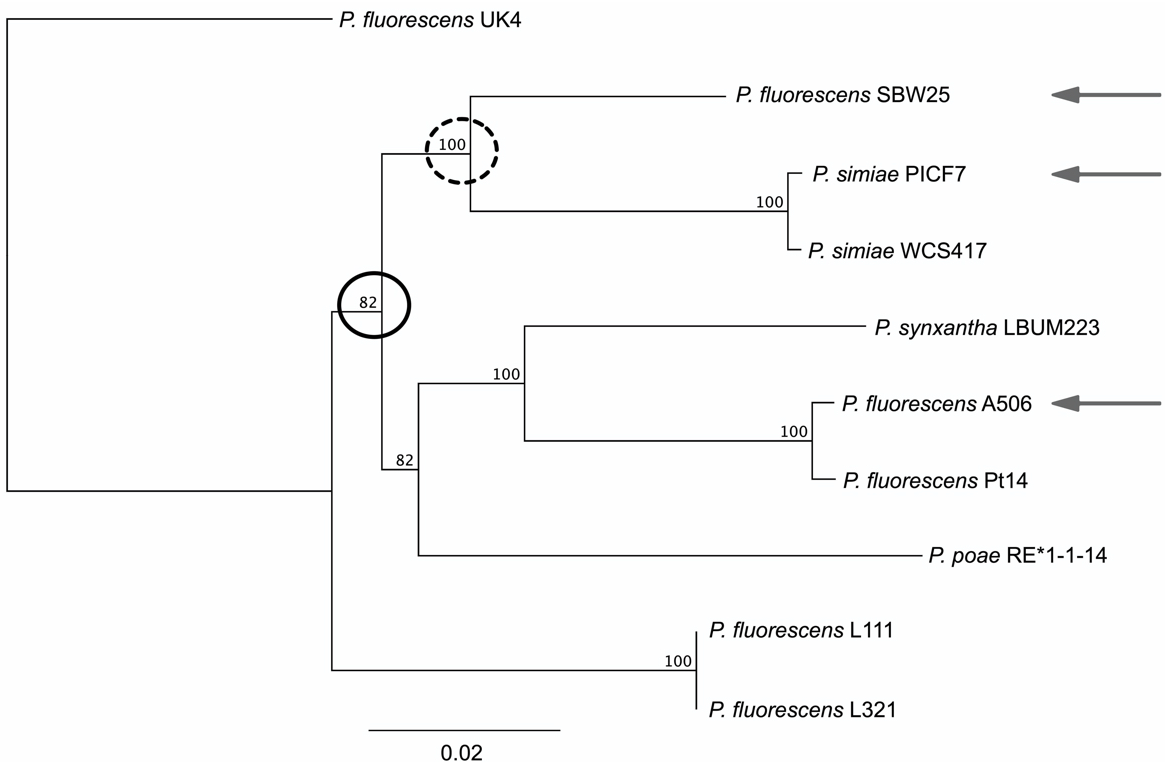
Phylogenetic tree highlighting the evolutionary relationships between *P. fluorescens* SBW25, *P. simiae* PICF7, and *P. fluorescens* A506. The three strains of interest are indicated by grey arrows. The last common ancestor of SBW25 and PICF7 is indicated by the dotted circle, and the last common ancestor of all three is indicated by a solid circle. The support shown next to the branches is based on 100 bootstrap replicates.

SBW25, PICF7 and A506 encode the biosynthetic genes required to synthesize distinct sets of EPSs; each strain carries genes for Pga, Psl, and alginate biosynthesis, while SBW25 and PICF7 encode genes for the biosynthesis of additional polymers (cellulose and colanic acid in SBW25, Pel in PICF7) (Table 1; Table S2). In addition, all three strains carry homologues of the *wsp*, *aws*, and *mwsR* loci, each of which is predicted to contain at least one c-di-GMP metabolizing domain (DGC/PDE; Table S2), Hence, SBW25, PICF7, and A506 each has the genetic capacity to over-produce c-di-GMP, with a strain-specific set of possible downstream (EPS) targets.

**Table 1.** A (non-exhaustive) list of genetic loci involved in EPS production in SBW25, PICF7, and A506. Further details of selected loci are provided in Table S2. Synonyms include *awsXRO/tpbB/yfiBNR*; *mwsR/morA*.

| Locus | Function | SBW25 | PICF7 | A506 |
| --- | --- | --- | --- | --- |
| *wss* | Cellulose biosynthesis | ✓ | ✗ | ✗ |
| *wca* | Colanic acid capsule biosynthesis | ✓ | ✗ | ✗ |
| *pel* | Pel biosynthesis | ✗ | ✓ | ✗ |
| *pga* | Pga biosynthesis | ✓ | ✓ | ✓ |
| *psl* | Psl biosynthesis | ✓ | ✓ | ✓ |
| *alg* | Alginate biosynthesis | ✓ | ✓ | ✓ |
| *wsp* | Regulation of EPS production | ✓ | ✓ | ✓ |
| *aws* | Regulation of EPS production | ✓ | ✓ | ✓ |
| *mwsR* | Regulation of EPS production | ✓ | ✓ | ✓ |

### PICF7 and A506 form emergent mats and diverse colony morphotypes in static microcosms

Mat formation by PICF7 and A506 was investigated relative to SBW25. Ten single (smooth, wild-type) colonies from each of the three strains were inoculated into separate microcosms, and incubated statically. By 72 hours, mats had emerged at the air-liquid interface of all microcosms. The mats formed from PICF7 were very similar in appearance to those formed from SBW25, while A506 mats were noticeably thinner, less defined, and usually less fluorescent (Figure 2A). Plating of cells from the 72-hour microcosms revealed the development of considerable colony diversity (Figure 2B). As expected from previous studies, SBW25 gave rise to three distinct morphotypes (Smooth, Wrinkly Spreader, Fuzzy Spreader) (Rainey and Travisano, 1998). Five distinct morphotype classes were observed arising from the majority of PICF7 microcosms (Smooth, Large Smooth, Wrinkly Spreader, Small Disc, Large Disc), while A506 microcosms typically gave rise to two morphotype classes (Smooth, Web). Notably, the PICF7 wrinkly spreaders (WS) closely resembled the mat forming WS morphotype of SBW25, and were seen arising from all ten microcosms. No wrinkly spreader colonies were observed arising from any of the A506 microcosms.

**Figure 2.**
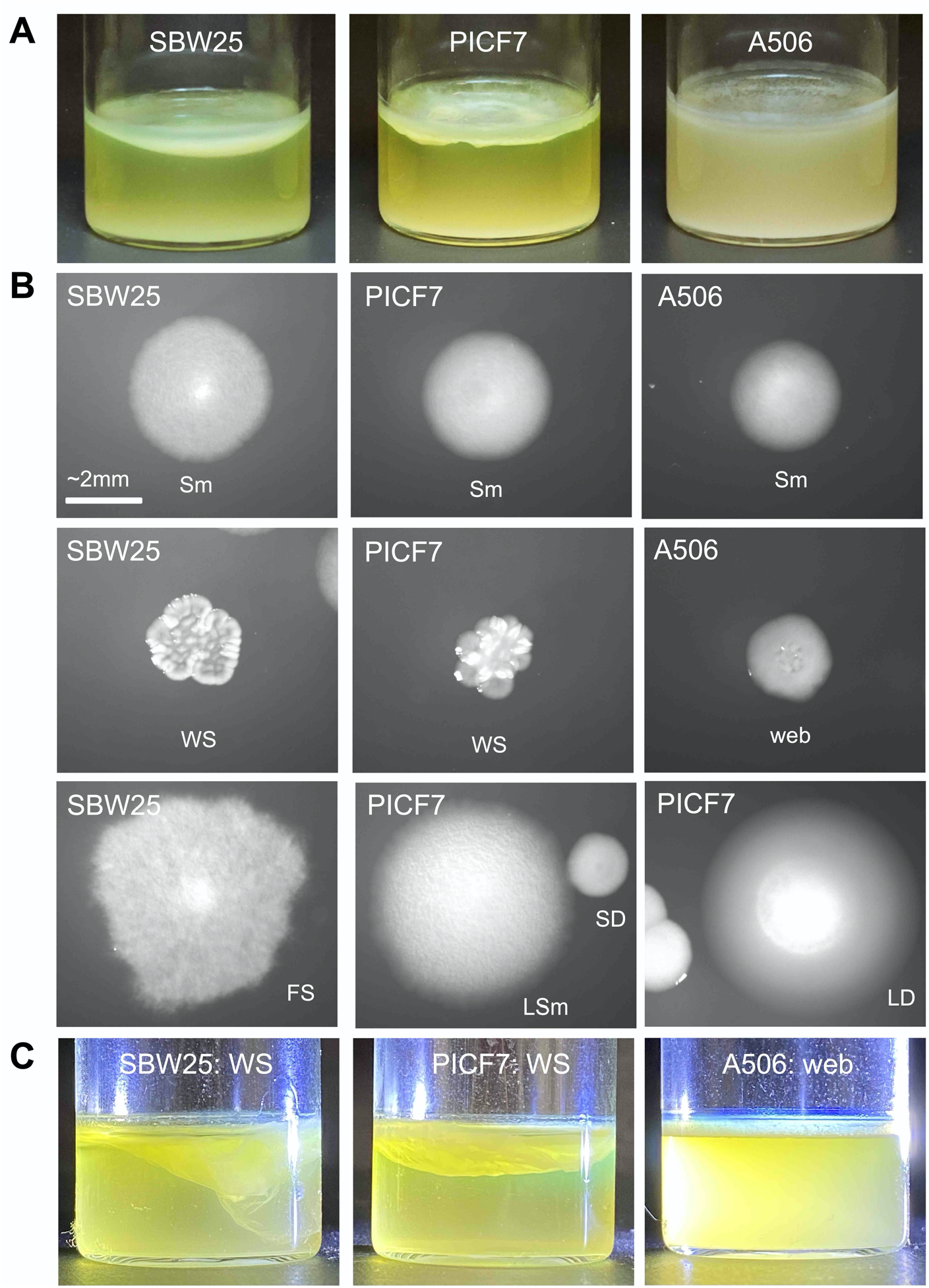
Mat formation by, and emergent colony diversity in, PICF7 and A506 relative to SBW25. (**A**) Representative 72-hour microcosms founded by SBW25, PICF7, and A506. All three strains develop mats at the air-liquid interface within 72 hours. (**B**) Dilution plating from microcosms shows ten morphotypes: three in SBW25 (Smooth (Sm), Wrinkly Spreader (WS), Fuzzy Spreader (FS)), five in PICF7 (Smooth (Sm), Wrinkly Spreader (WS), Large Smooth (LSm), Large Disc (LD), and Small Disc (SD)), and two in A506 (Smooth (Sm), and Web). (**C**) SBW25-WS, PICF7-WS, and A506-Web give rise to mats at the air-liquid interface within 24 hours. See Figure S1 for 24-h microcosms from other morphotypes. Brightness and exposure of some images were altered in Preview.

While PICF7 and A506 form mats in 72-h microcosms, the mats typically require between 48 and 72 hours to fully emerge. To test which, if any, of the emergent colony morphotypes gained the ability to form stable mats more quickly, examples of each morphotype were purified and inoculated into fresh microcosms. After 24 hours, five of the ten morphotypes formed a stable mat: SBW25 wrinkly spreader (SBW25-WS), PICF7 wrinkly spreader (PICF7-WS), PICF7 large smooth (PICF7-LSm), PICF7 small disc (PICF7-SD), and A506 web (A506-Web) (Figure 2C, Figure S1).

The results in this section demonstrate that both PICF7 and A506 develop mats in 72-h microcosms, and that mat formation is accompanied by the emergence of a range of colony morphotypes. In PICF7, at least three morphotypes may contribute to the 72-h mat (WS, LSm, SD), while the A506 mat is likely underpinned by the web morphotype. In the remainder of this work, we concentrate on characterizing the molecular mechanisms underpinning mat formation in the emergent PICF7-WS and A506-Web morphotypes.

### *PICF7 wrinkly spreaders carry mutations in* wsp, aws, or mwsR

The mats and WS colonies observed from PICF7 static microcosms strongly resemble their SBW25 counterparts. That is, they have the emergent property of forming a stable mat within 24 hours, and the wrinkly phenotype is heritably stable (*i.e.*, when PICF7-WS colonies are sub-streaked on agar, they give rise to exclusively WS colonies). In order to determine whether the same mutational pathways underpin the WS phenotype in PICF7 as SBW25, ten WSs were isolated from independent, 72-h microcosms founded by PICF7 (giving PICF7-WS1 to 10) and SBW25 (giving SBW25-WS1 to 10). Whole genome sequencing of these twenty WS genotypes revealed a mutation in *wsp, aws*, or *mwsR* in every case (Table 2; Text S1). Six PICF7-WS types carried mutations in the *wsp* locus (versus four SBW25-WSs), three in the *aws* locus (versus two SBW25-WSs), and one in *mwsR* (versus four SBW25-WSs). Each mutation is predicted to result in c-di-GMP over-production by WspR, AwsR, or MwsR, leading to the activation of EPS biosynthesis and mat formation (Text S1).

**Table 2.**
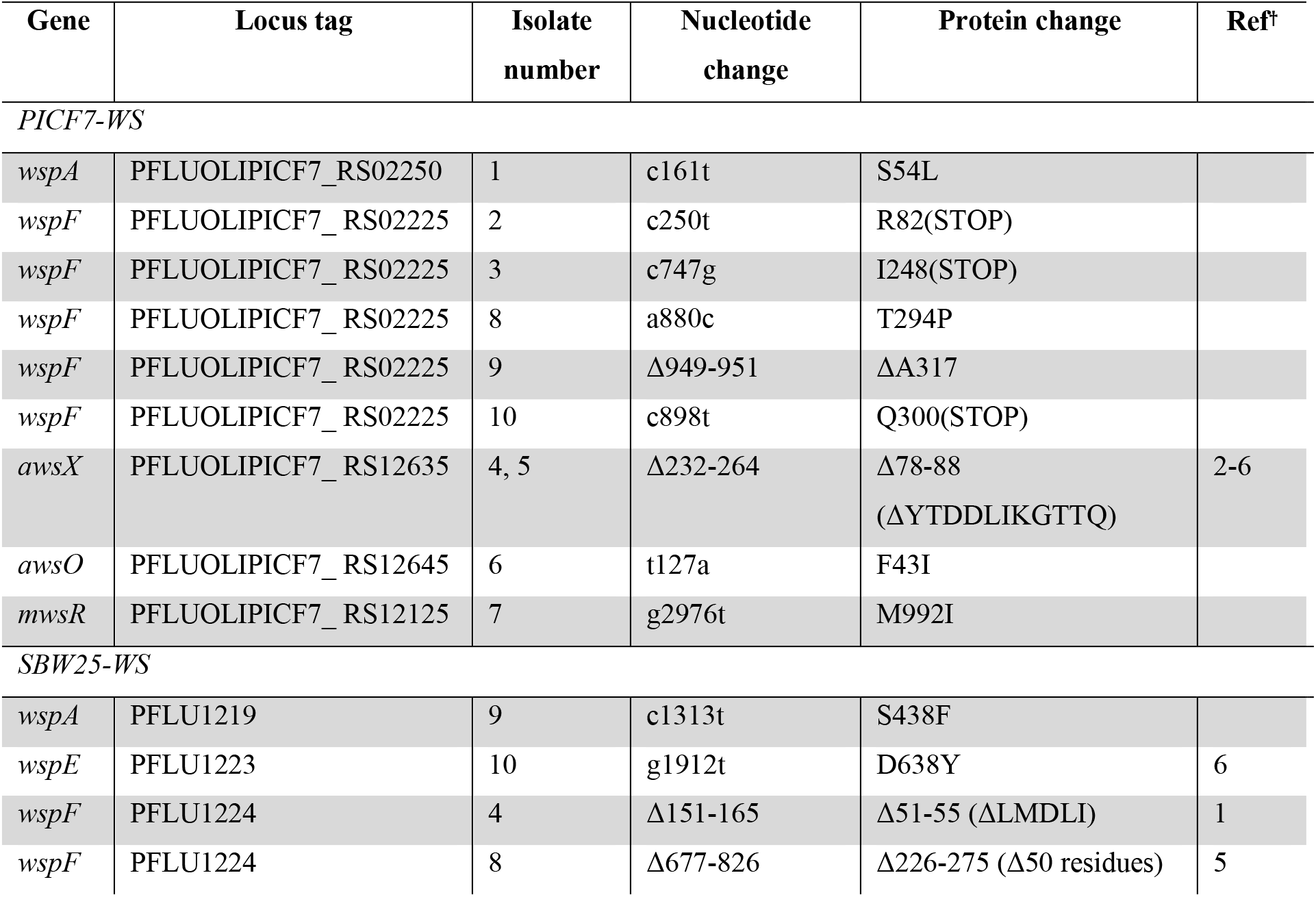
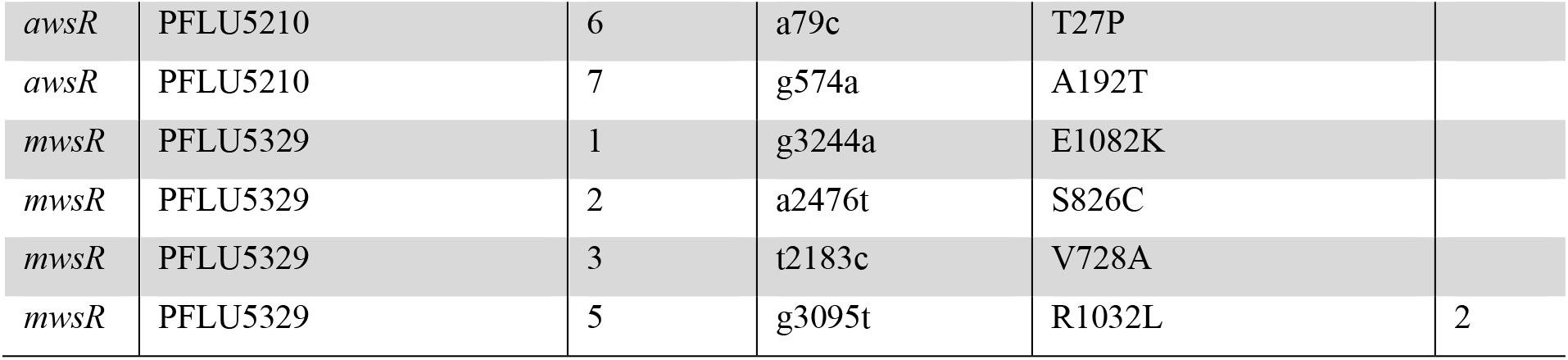
Details of mutations identified in ten WS strains isolated from each of PICF7 and SBW25. **†**Reference is provided if a change in the same amino acid(s) has previously been reported to cause the WS phenotype in *P. fluorescens* SBW25. 1=Bantinaki *et al.*, 2007, 2=Gallie *et al.*, 2015, 3=Gallie *et al.*, 2019, 4=Lind *et al.*, 2017, 5=Lind *et al.*, 2019, 6=McDonald *et al.*, 2009. See Text S1 for further molecular details.

### The Pel polymer contributes to the structure of the PICF7 wrinkly spreader phenotype

Despite the phenotypic and genotypic similarities between PICF7-WS and SBW25-WS, the mats formed by each must differ on a structural level because PICF7 lacks homologues of the cellulose-biosynthetic *wss* genes that are activated by *wsp, aws,* or *mwsR* mutations in SBW25-WS. However, PICF7 carries several other polymer biosynthetic loci that could conceivably be activated by *wsp, aws,* or *mwsR* mutations (see Table 1). To investigate the structural basis of the PICF7-WS phenotype, a suppressor analysis was performed on PICF7-WS3, -WS4, -WS7, and -WS8 (carrying mutations in *wspF*, *awsX*, *mwsR*, and *wspF*, respectively). Each genotype was subjected to random transposon mutagenesis, and transposon mutants that had reverted to the smooth colony phenotype, and lost the ability to form 24-h mats in static microcosms, were obtained. The genomic location of the transposon was determined in these mutants. Overall, approximately 15,600 transposon mutants were screened, and the insertion site determined in 100 suppressor mutants (Table S4).

Multiple independent transposon insertions were obtained in two types of loci. Firstly, 29 independent insertions were obtained in the *wsp, aws,* or *mwsR* loci. Presumably, these insertions directly counteract the WS mutation in each background, lowering c-di-GMP levels and reversing the WS phenotype. Secondly, 14 independent insertions were obtained in the seven-gene EPS biosynthetic locus, *pelA-G* (PFLUOLIPICF7_RS06785-PFLUOLIPICF7_RS06755; Figure 3A). It is probable that this class of suppressor mutants maintains high c-di-GMP levels, but loses the WS phenotype because the downstream EPS target (Pel) is inactivated. In three *pel* transposon mutants – carrying insertions in *pelA* (PFLUOLIPICF7_RS06785), *pelD* (PFLUOLIPICF7_RS06770), and *pelG* (PFLUOLIPICF7_RS06755) – the majority of the transposon was removed by Cre-mediated recombination, eliminating possible polar effects. The Cre-deletion genotypes gave smooth colonies and were unable to form 24-h mats (Figure 3B-C), strongly indicating that at least three of the seven *pel* genes are essential for the PICF7-WS phenotype. We conclude that Pel is an important, c-di-GMP regulated, structural component of the PICF7-WS mat.

**Figure 3.**
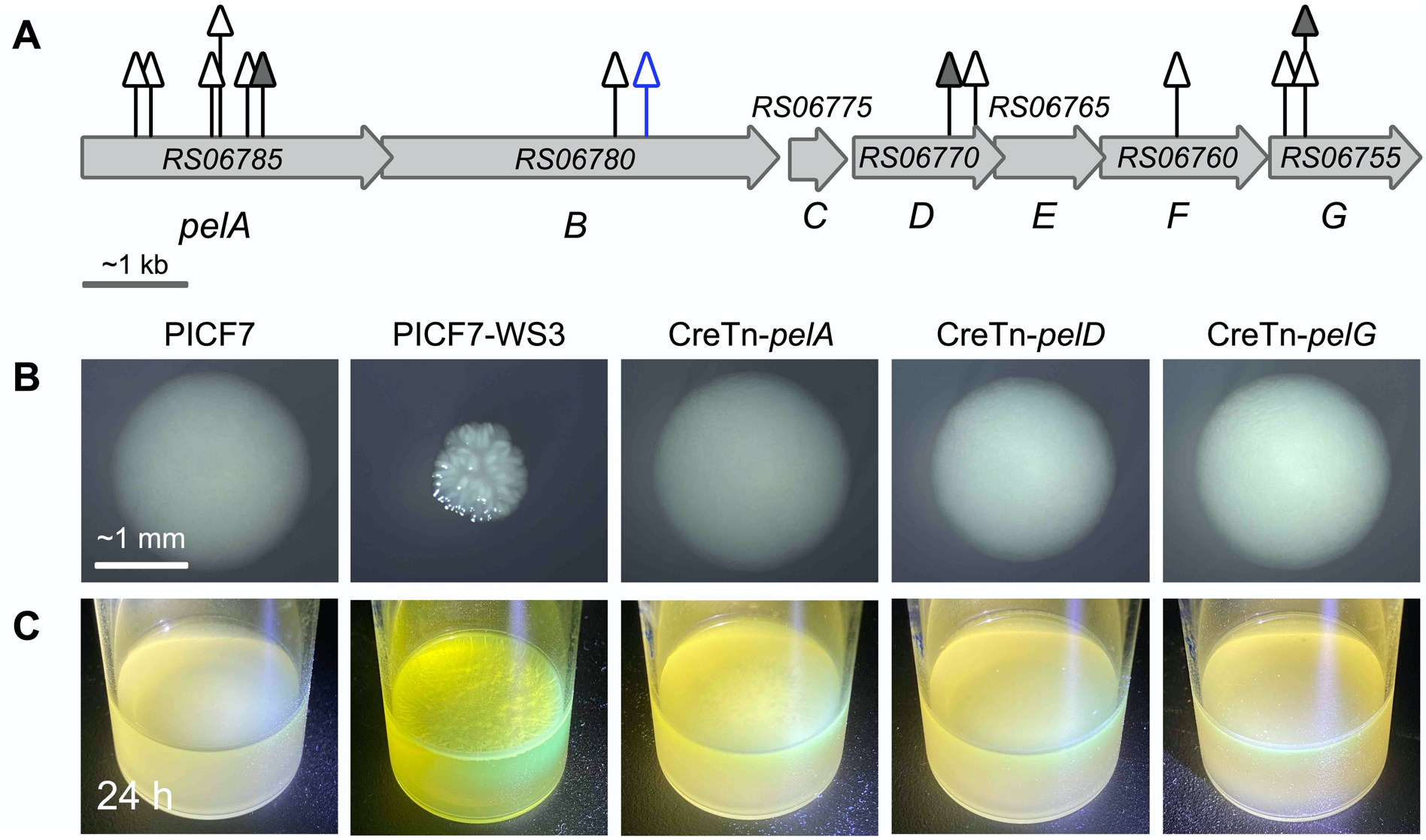
Pel production is a major structural component of the PICF7-WS phenotype. (**A**) Fourteen independent WS suppressors were obtained in five of the seven genes of the PICF7 *pel* locus (*pelA-G*; Tables S2 and S4). Grey triangles=transposon mutants obtained in the background of a *wspF* mutation (PICF7-WS3, PICF7-WS8); blue triangle=transposon mutant obtained in the background of the *awsX* mutation (PICF7-WS4); open triangles=mutants carrying the full transposon; solid triangles=mutants in which a Cre-mediated deletion of the original transposon has been constructed (giving CreTn-*pelA*, CreTn-*pelD*, CreTn-*pelG*). The Cre-deleted transposon mutants have lost the WS colony morphology on KB agar (**B**), and no longer form strong mats in static microcosms within 24 hours (**C**). Brightness and exposure of some images were altered in Preview.

### The A506-Web colony phenotype is semi-heritable

In contrast to SBW25-WS and PICF7-WS, the A506-Web phenotype is only semi-heritable. Firstly, A506-Web colonies derived from 72-hour static microcosms rapidly and repeatedly give rise to a mixture of web and smooth (*i.e.*, wild type-like) colonies when plated on KB agar (Figure 4A). Secondly, when A506-Web colonies were grown in shaken overnight culture (*i.e.*, an environment that precludes mat formation) and plated on KB agar, between 9 % and 45 % of colonies reverted to the ancestral, smooth morphotype (Figure 4B). No such colony diversity was observed arising from the colonies of the stable mat-forming morphotypes (A506-*wspF*-LoF, SBW25-WS, PICF7-WS, PICF7-LSm, or PICF7-SD) (Figure 4B). Of the web and smooth colony types arising from A506-Web, only the web type retained the ability to give rise to 24-h mats in static microcosms (Figure 4C).

**Figure 4.**
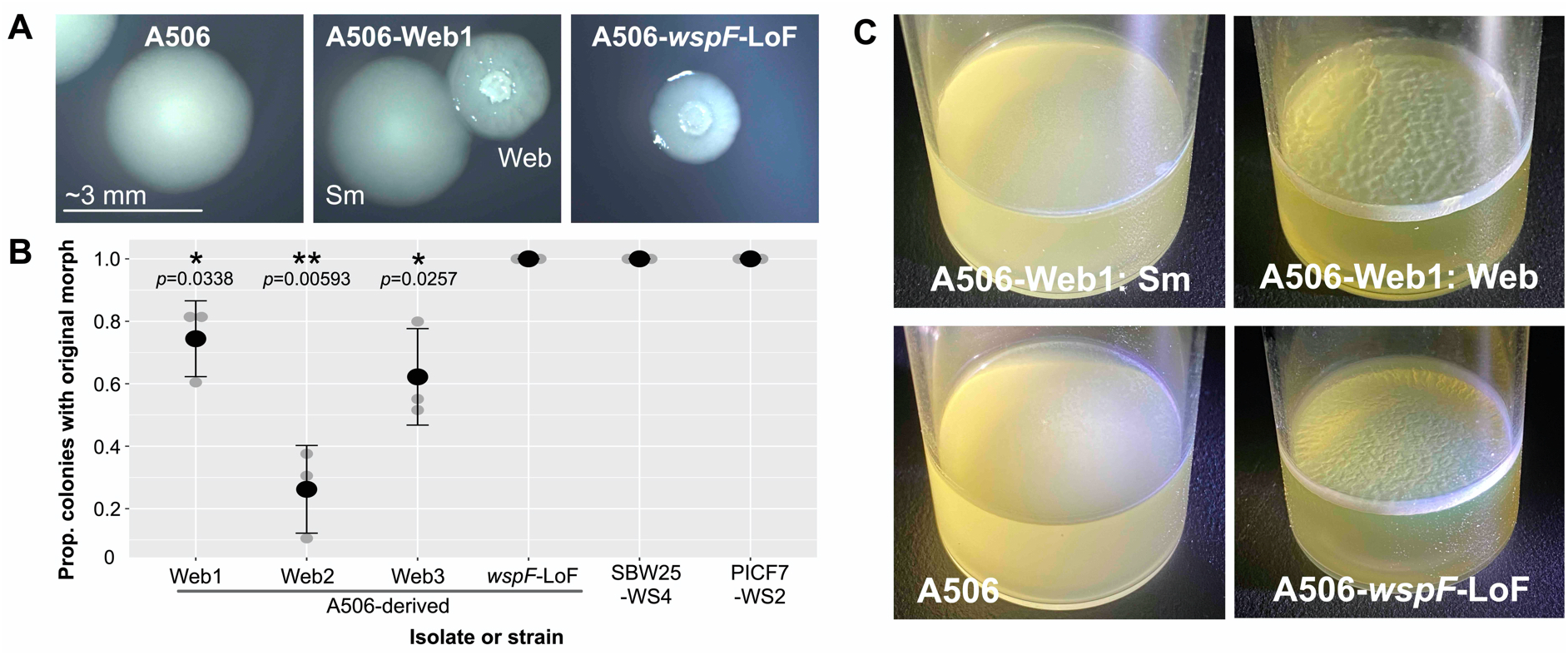
Semi-heritable (A506-Web) and heritable (A506-*wspF*-LoF) routes to mat formation in A506. (**A**) A506-Web isolates from 72-h static microcosms give rise to a mixture of smooth (Sm) and web colonies on KB agar (A506-Web1 is pictured as an example). A506-*wspF*-LoF, carrying a loss-of-function mutation in *wspF*, consistently generates colonies with a similar, web-like appearance. (**B**) Quantification of colony morphotype loss demonstrated significant degree of web morphotype loss in A506-Web colonies (one-tailed, one sample *t*-test *p*<0.0338), no loss of the web-like morphotype in A506-*wspF*-LoF, and no loss of WS morphotype in SBW25-WS4 or PICF7-WS2 (both carrying loss-of-function *wspF* mutations). Three biological replicates were performed for each strain, with a minimum of 45 colonies counted across three plates *per* replicate. Grey dots are individual data points, black dots are means, and black whiskers are standard errors. (**C**) Like A506, smooth colonies isolated from A506-Web1 in panel A (A506-Web1:Sm) do not give rise to a mat in 24-h static microcosms. Both the web colony counterparts (A506-Web1:Web) and A506-*wspF*-LoF form a strong mat within 24 hours. Brightness and exposure of some images were altered in Preview.

The relative ease with which the A506-Web phenotype reverts to the wild-type morphotype suggests that, unlike PICF7-WS and SBW25-WS, it is not underpinned by a stable genetic change. Indeed, no SNPs or indels were confirmed on the A506 chromosome or plasmid during whole genome re-sequencing of the ten independent A506-Web isolates (Table S3). Interestingly, evidence of large-scale, tandem duplication events was found in all ten A506-Web isolates, suggesting a flexible region of the chromosome to one side of the replication terminus (Text S2). The affected regions vary in size (~158 kb to ~634 kb) and precise location (between genomic positions 1,900,000 bp and 2,897,000 bp), but contain a shared segment of ~126 kb beginning at genomic position ~2,314,000 bp. While intriguing, these amplifications are not thought to contribute to the web phenotype, because (i) a similar region was observed upon sequencing the A506 wild-type, which had not been subjected to growth in a static microcosm, and (ii) the set of ~115 genes encoded in the shared region does not include any genes with known roles in polymer synthesis or regulation (*e.g.*, *pga, psl, wsp, aws, mwsR*).

### *A deletion in* wspF *leads to a stable, web-like phenotype in A506*

Even though *wsp*/*aws*/*mwsR* mutations were not identified in A506-Web isolates, seemingly complete homologues of all three loci exist (Table 1; Table S2). This begs the question of whether mutations in these loci *could* generate differences in colony morphology, and 24-h mats. To investigate, a scar-free, loss-of-function mutation was constructed in the A506 *wspF* gene (Δ151-165 in PflA506_1189, leading to the deletion of amino acids 51-55; see Text S3). This mutation is a commonly occurring loss-of-function mutation underpinning the SBW25-WS phenotype (Bantinaki *et al.*, 2007; Gallie *et al.*, 2019), and was also identified in SBW25-WS4 of this study. The resulting genotype, A506-*wspF-* LoF, forms colonies that are similar in appearance to A506-Web colonies (“web-like”), and gives rise to 24-h mats (Figure 4A, 4C). Further, the web-like colony phenotype is stable; A506-*wspF*-LoF web-like colonies give rise to exclusively web-like colonies when streaked on agar plates, and when plated after growth in shaken overnight culture (Figure 4B). Hence, A506 carries the genetic and molecular machinery required to produce mats by at least one of the same mutational routes as SBW25-WS and PICF7-WS (loss-of-function mutations in *wspF*). However, the emergence of mat formation by a readily reversible route appears to be favoured in our experiments.

### Pga and Psl contribute to A506 mat formation

Molecular routes to mat formation by A506 are expected to culminate in the over-production of EPS (and/or other surface components). There are at least three candidate EPSs: Pga, Psl, and alginate (Table 1). Two of these, Pga and Psl, have been shown to be regulated by c-di-GMP in other pseudomonads (Borlee *et al.*, 2010; Malone *et al.*, 2010; Lind *et al.*, 2017). We initially focused on Pga, because A506-Web colonies share morphological similarities with the SBW25 <u>P</u>ga <u>w</u>rinkly <u>s</u>preader (PWS). PWSs are WS-like genotypes, often carrying *aws* or *mwsR* mutations, isolated from microcosms inoculated with an SBW25 strain deficient in cellulose biosynthesis (SBW25-Δ*wss*) (Lind *et al.*, 2017).

To test the possibility that Pga plays a role in heritable mat formation, the four-gene *pga* operon (*pgaA-D;* PflA506_0154 to PflA506_0157) was cleanly deleted from A506-*wspF-*LoF, and the effect on colony and mat forming phenotypes assessed (Figure 5A). Elimination of Pga noticeably altered heritable mat formation; A506-*wspF*-LoF-Δ*pga* formed visibly weaker 24-h mats than A506-*wspF*-LoF, and 72-h mats with a different texture. Deletion of *pga* from A506 had no immediately obvious phenotypic effects (Figure 5B), suggesting that while Pga may play a structural role in A506 mat formation, other EPSs can fulfill this role if required (perhaps in a similar manner to Pga substituting for cellulose in mats derived from SBW25-Δ*wss*).

**Figure 5.**
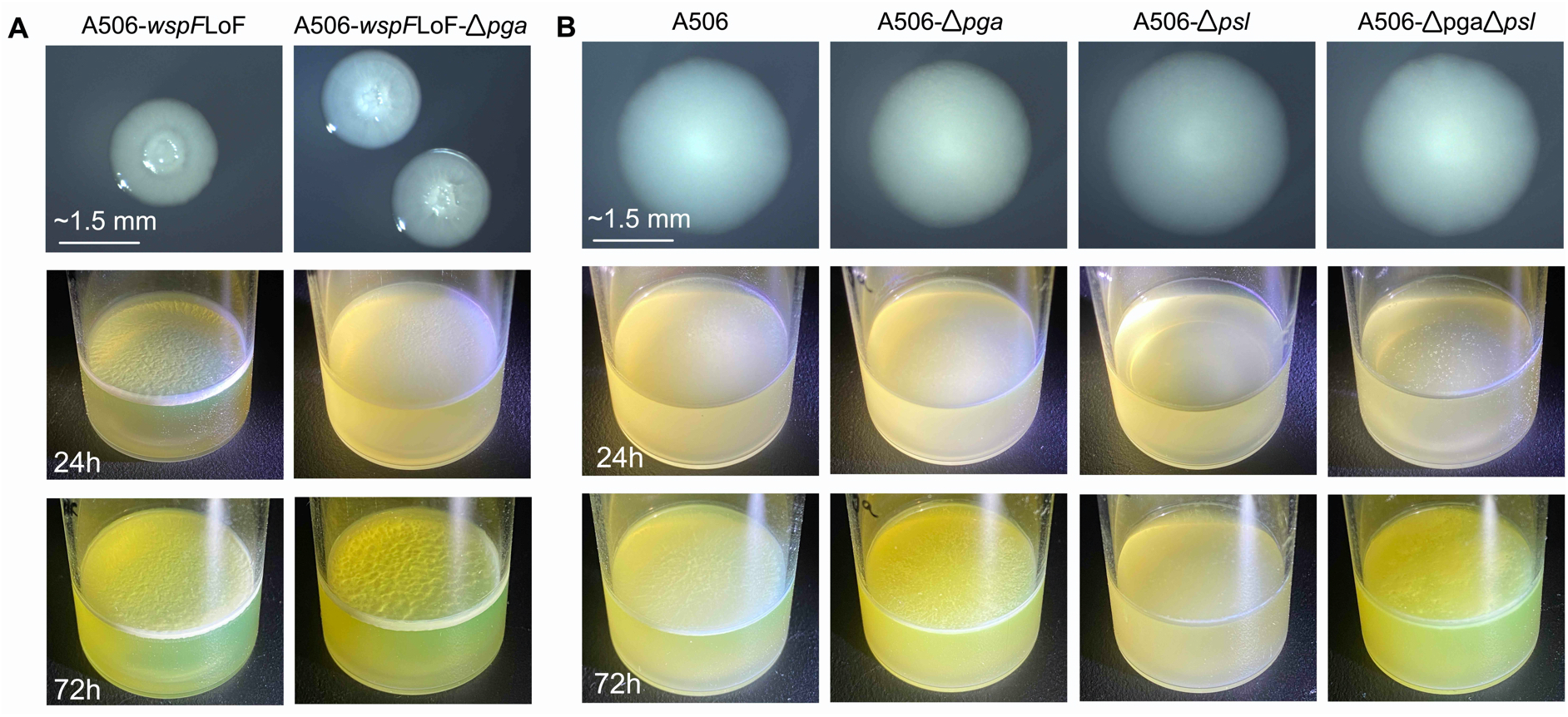
Pga and Psl contribute to A506 mat formation. (**A**) The effect of scar-free deletion of the *pga* operon from A506-*wspF*-LoF on colony morphology (top), 24-h mat formation (middle), and 72-hour mat formation (bottom). (**B**) Same as panel A, but with *pga, psl*, or both deleted from A506. All photographs within a single row were taken under the same conditions, at the same time point. Brightness and exposure of some images were altered in Preview.

The role of various structural components in mat formation by the emergent A506-Web type is not easy to elucidate, due to the semi-heritable nature of the phenotype; the phenotype is quickly lost during the successive rounds of growth required for the transposon mutagenesis and/or the genetic engineering protocol. Therefore, we deleted the *pga* (*pgaA-D;* PflA506_0154 to PflA506_0157) and *psl* (*pslA-K;* PflA506_1982 to PflA506_1971) loci, separately and in combination, from wild-type A506 (giving A506-Δ*pga*, A506-Δ*psl*, and A506-Δ*pgaΔpsl*). Elimination of both *pga* and *psl* altered, but did not entirely eliminate, the emergence of 72-h mats from A506 (Figure 5B). Overall, the results in this section indicate that while Pga and Psl play a role in mat formation, other structural components may also contribute to the emergence of A506 mats.

## DISCUSSION

*P. fluorescens* SBW25 readily acquires mutations in *wsp, aws,* or *mwsR* that elevate c-di-GMP, leading to the constitutive synthesis of cellulose and the emergence of mats at the air-liquid interface. Here, we have investigated the divergent mechanisms by which SBW25 relatives *P. simiae* PICF7 and *P. fluorescens* A506 produce similar, emergent mats. The details of mat formation in PICF7 and A506 are discussed below, and a summary model is provided in Figure 6.

**Figure 6.**
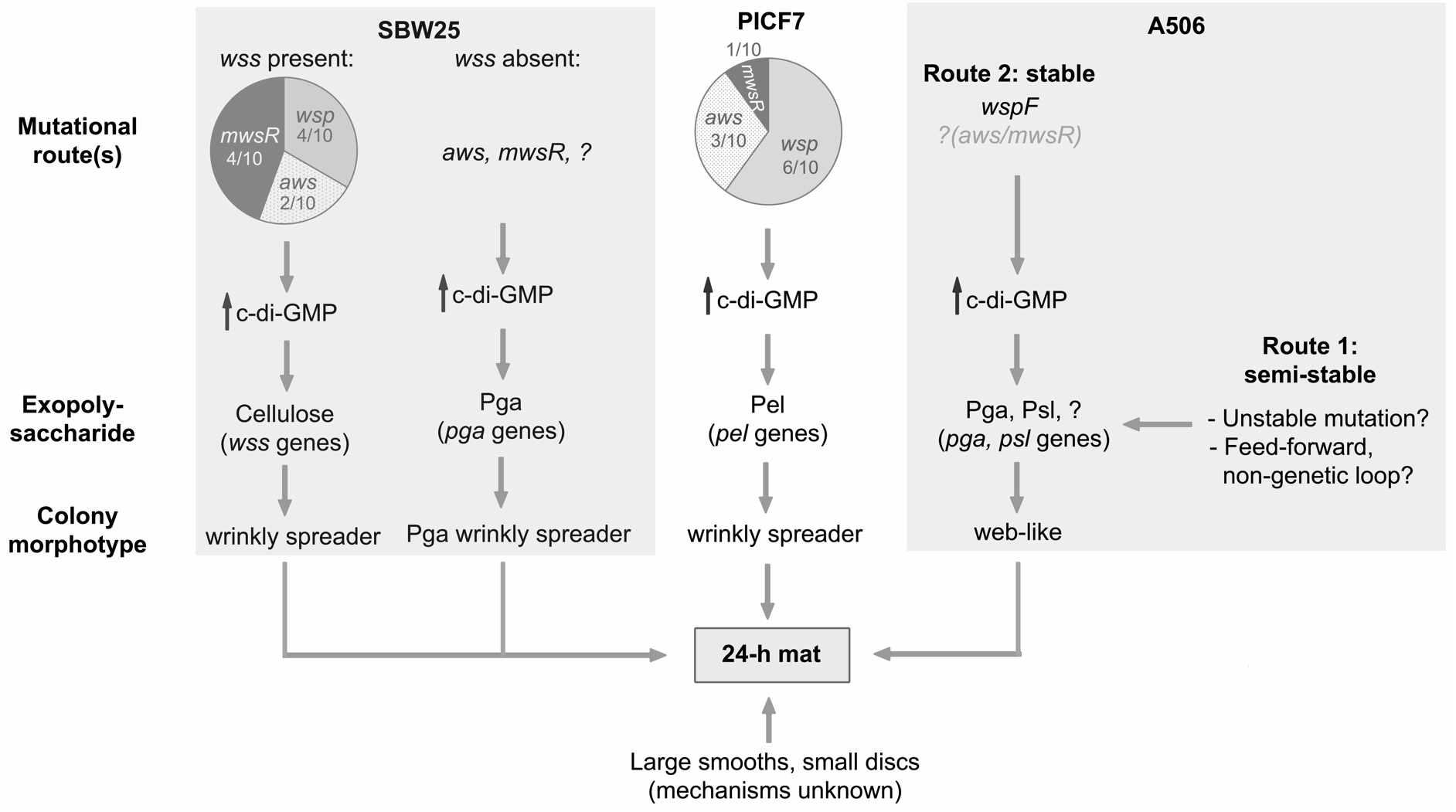
Model of the molecular routes to 72-h mat formation in *P. simiae* PICF7 and *P. fluorescens* A506, as compared with the previously well-studied *P. fluorescens* SBW25. Pie charts were drawn using mutations identified by genome sequencing of ten SBW25 and ten PICF7 WS genotypes (see Table 2).

### A model for mat formation by PICF7-WS

One morphotype that contributes to the emergence of mats from PICF7 is the wrinkly spreader (PICF7-WS). There are notable parallels between PICF7-WS and SBW25-WS. Namely, (i) they produce similar-looking, heritably stable, wrinkly colonies and 24-h mats, and (ii) these phenotypes are underpinned by c-di-GMP-elevating mutations in *wsp*, *aws*, or *mwsR* homologues. However, the downstream targets of c-di-GMP differ in the two strains. In SBW25, elevated c-di-GMP levels stimulate the mass production of cellulose by the Wss biosynthetic machinery (Spiers *et al.*, 2002; Gal *et al.*, 2003; Goymer *et al.*, 2006). PICF7 lacks *wss* homologues, and cannot synthesize cellulose. Instead, the PICF7-WS phenotype requires the over-expression of Pel (Figure 6).

Pel is a cationic, glucose-rich EPS that has mainly been characterized as a structural component of *P. aeruginosa* biofilms (Jennings *et al.*, 2015). In *P. aeruginosa*, c-di-GMP-mediated over-production of Pel is required for the emergence of WS-like, rugose <u>s</u>mall-<u>c</u>olony <u>v</u>ariants (RSCVs) (D’Argenio *et al.*, 2002; Hickman *et al.*, 2005; Starkey *et al.*, 2009). C-di-GMP-mediated regulation of *P. aeruginosa* Pel production occurs through FleQ, a c-di-GMP binding transcriptional regulator. FleQ binds to two stretches of DNA upstream of *pelA*, and, in the absence of c-di-GMP, represses *pel* transcription (Hickman and Harwood, 2008). When DNA-bound FleQ interacts with c-di-GMP, the conformation of the region changes such that FleQ becomes a transcriptional activator of *pel* (Baraquet *et al.*, 2012; Matsuyama *et al.*, 2016). FleQ is also involved in c-di-GMP dependent EPS production in *Pseudomonas putida* (Ramos-González *et al.*, 2016; Molina-Henares *et al.*, 2017).

The above suggests that FleQ may play a role in c-di-GMP mediated regulation of Pel production in PICF7. Indeed, a recent study reported that a PICF7 strain carrying an inactivating insertion in *fleQ* (PFLUOLIPICF7_16515) resulted in altered capacity to form a biofilm (Montes-Osuna *et al.*, 2021). The extent to which, and mechanisms whereby, *fleQ* and other regulators are involved in PICF7 Pel regulation remain to be elucidated.

### Additional routes to mat formation in PICF7

In addition to WS, 72-h PICF7 microcosms were typically observed to contain multiple other emergent colony morphotypes (LSm, SD, LD). Each morphotype has its own distinguishing phenotypic features, which presumably result from different molecular processes. With the exception of WS, these molecular processes remain almost entirely uncharacterized; they may or may not involve c-di-GMP, and may each regulate the expression of different downstream EPSs or cell surface components. We do know that at least three of the emergent morphotype classes – WS, LSm and SD – are heritable (*i.e.*, are likely underpinned by stable mutations). In addition, these three morphotypes can each independently form 24-h mats (see Figure S1), suggesting that each type arises and competes for dominance in the emerging, 72-h mat. The relative proportions of each morphotype have not yet been studied, but these are likely to depend on a combination of the rate at which each morphotype arises (*e.g.*, mutational target size and localized mutation rates), and the relative fitness of each type in the emerging population.

The co-emergence of multiple, distinct mat-forming morphotypes strongly suggests that 72-h PICF7 mats consist of phenotypically and genotypically heterogeneous subpopulations, possibly with each morphotype producing distinct EPSs, and/or cell surface components. PICF7 has the genetic capacity to produce several EPSs (including Pel, Pga, Psl, and alginate), and various cell surface components (*e.g.*, adhesins, lipopolysaccharides, pili). Expression of each of these components is expected to confer specific physicochemical properties on the surrounding cells (reviewed in Rehm, 2010), some of which may be beneficial in the natural plant environment.

### Molecular routes to semi-heritable mat formation by A506-Web

The emergence of A506 mats over 72 hours is underpinned by the semi-heritable A506-Web morphotype, which forms colonies with a web-like internal structure that can form full mats within 24 hours. The A506-Web phenotype is less stable than the mat-forming morphotypes arising from PICF7 and SBW25. The degree of morphotype stability provides some insight into the underlying molecular causes of the phenotype. At one end of the scale, a readily reversible phenotype is likely to be underpinned by non-heritable mechanisms (*e.g.*, activation of an EPS transcription factor in the presence of a particular environmental stimulus; see introduction). Such mechanisms can be switched on and off within the lifetime of a cell, and are thus not heritable. At the opposite end of the scale, an (almost) irreversible phenotype is likely to be underpinned by a stable genetic change (*e.g.*, a deletion in *wspF* giving rise to the WS phenotype). In the absence of further mutation(s), the resulting phenotype is stable and heritable. The semi-heritable A506-Web phenotype is not expected to result from either a straightforward transcriptional change (no heritability), or a stable genetic change (heritability), but a mechanism that generates semi-heritability.

We see two hypothetical possibilities for a mechanism underpinning semi-heritability (Figure 6). Firstly, the A506-Web phenotype could conceivably result from a non-stable genetic change that flips between two states at relatively high frequency (*i.e.*, phase variation). These types of mutations – examples of which include slippage in homopolymeric tracts (De Bolle *et al.*, 2000; Orsi *et al.*, 2010), site-specific inversions (Abraham *et al.*, 1985; Dybvig and Yu, 1994), and tandem duplication events (Anderson and Roth, 1981; Ayan *et al.*, 2020) – occur and revert randomly, but at frequencies several orders of magnitude higher than standard SNPs (reviewed in Anderson and Roth, 1977; Moxon *et al.*, 2006). If such a high-frequency mutational locus were to exist within, for example, an A506 EPS biosynthetic locus, an initially clonal, growing population would rapidly and repeatedly generate sub-populations of “EPS on” (web) and “EPS off” (wild-type). Once both forms exist, selection would act to influence the relative proportion of each form, with web and wild-type dominating in static and shaken environments, respectively. While we did not find evidence of mutable loci underpinning the unstable A506-Web phenotype, we note that many kinds of high-frequency mutations are notoriously difficult to identify using whole genome re-sequencing. However, the rate at which the A506-Web morphotype repeatedly reverted to the wild-type morphotype in overnight shaken culture was observed to be much higher than would typically be expected for phase variants (Anderson and Roth, 1977; Moxon *et al.*, 2006).

A second possibility is that no genetic change is required to switch between A506-Web and wild-type morphotypes. Instead, semi-heritability may be achieved by a molecular network topology that can generate two distinct, semi-stable phenotypic states (“EPS on” or “EPS off”) (Tiwari *et al.*, 2011). To illustrate, imagine the following example. In the static microcosm, a signal activates expression of a transcription factor (TF), which in turn activates the transcription of (i) a DGC and/or EPS biosynthetic genes, and (ii) itself. This leads to (i) the “EPS on” state and mat formation, and (ii) self-perpetuating TF production that can persist even after the initial environmental stimulus is removed. A switch back to the “EPS off” state requires TF levels to stochastically drop below a particular threshold, perhaps as a result of stochastic division of intracellular protein content upon cell division. Examples of bistable molecular networks have been shown to underpin semi-heritable, ON/OFF expression of colanic acid-like capsules in *P. fluorescens* SBW25 (Gallie *et al.*, 2015; Gallie *et al.*, 2019; Remigi *et al.*, 2019), bistable expression of the PDE PdeL in *E. coli* (Reinders *et al.*, 2016), and bistable white-opaque switching in *Candida albicans* (Zordan *et al.*, 2006). Some bistable switches can switch between states more rapidly than is achieved by phase variation (*e.g.*, Gallie *et al.*, 2015; Remigi *et al.*, 2019), and hence could conceivably be involved in the A506-Web phenotype.

The existence of a semi-heritable mechanism underpinning A506 mat formation provides a degree of phenotypic flexibility for A506 that has – at least so far – not been observed in laboratory populations of SBW25 and PICF7. Such a mechanism may enable A506 to readily and repeatedly switch between planktonic and sessile lifestyles, without the accumulation of irreversible, loss-of-function mutations in EPS biosynthetic pathways.

### A stable route to A506 mat formation: wspF mutations

In addition to the semi-heritable route, A506 can form stable web-like colonies and mats through at least one of the three mutational routes that underpin the SBW25-WS and PICF7-WS phenotypes: loss-of-function mutations in *wspF*, a negative regulator of the WspR DGC (Figure 6). Protein domain predictions for the Wsp pathway (see Table S2) are consistent with *wspF* mutations having similar downstream effects in all three strains: de-repression of WspR DGC activity, leading to an increase in c-di-GMP, and overproduction of at least one EPS (cellulose in SBW25, Pel in PICF7, and Pga in A506). Wsp-mediated control of these three distinct EPSs demonstrates a certain degree of evolutionary flexibility in mat structural components, indicating that EPS biosynthetic loci can be lost or gained, and may be readily incorporated into existing host regulatory networks.

It is currently unclear to what extent the web-like phenotype generated by mutation of *wspF* resembles the semi-heritable A506-Web phenotype (at both molecular and structural levels). However, while many mutational routes to mat formation are presumably available in A506 (*i.e.*, loss of function mutations in *wspF*), our results indicate that genotypes carrying these mutations do not dominate in emergent mats; all ten of our independently-isolated A506-Web types showed the semi-heritable web-like phenotype, and no *wspF* (or other) mutations were identified. The bias towards the semi-heritable route to mat formation may result from a higher rate of A506-Web generation than *wspF* mutation, and/or a selective advantage of A506-Web over *wspF* mutants.

The diversity of molecular and structural routes to mat formation in *P. fluorescens* SBW25, *P. simiae* PICF7, and *P. fluorescens* A506 demonstrates a high degree of evolutionary flexibility in EPS regulation, highlighting the ecological and evolutionary importance of mat and biofilm formation.

## EXPERIMENTAL PROCEDURES

### Phylogenetic tree construction

The complete genome sequences of ten close *Pseudomonas* strains were downloaded from NCBI in February 2021 (for reference numbers see Table S1). The most divergent strain, *Pseudomonas fluorescens* UK4, was used as an outgroup. A sequence alignment was generated using REALPHY version 1.13 (Bertels *et al.*, 2014). In REALPHY, three reference alignments were generated (A506, SBW25, and RE*-1-1-14) and merged. The merged alignment consisted of 1,464,229 alignment columns. From this alignment, a maximum likelihood tree was generated with PHYML (GTR substitution model) (Guindon and Gascuel, 2003), and visualized in Geneious (v11.1.4).

### Strains, plasmids, and primers

Details of all strains, plasmids, and oligonucleotides used in this study are provided in Table S1.

### Growth conditions

*E. coli* cloning strains were grown in Lysogeny Broth (LB) at 37°C (~18 hours, shaking). When not grown in static microcosms (see below), *Pseudomonas* strains were grown in King’s Medium B (KB; King *et al.*, 1954) at 28°C (~18 hours, shaking). *Pseudomonas* cells were plated on KB containing 1.5 % agar, and incubated at 28°C for 48 hours. Where appropriate, media were supplemented with: nitrofurantoin (NF; 100 μg ml^−1^), kanamycin (Km; 50 μg ml^−1^), tetracycline (Tc; 50 μg ml^−1^), and/or 5-Bromo-4-Chloro-3-Indolyl ß-D-Galactopyranoside (X-gal; 60 μg ml^−1^).

### Mat formation in static microcosms

Static microcosms were grown in 30 ml glass tubes containing 6 ml KB. Each microcosm was inoculated with 6 μl of stationary phase *Pseudomonas* culture, vortexed for 5 seconds, the lid loosened, and grown without agitation at 28°C for 24 or 72 hours. For analysis of emergent colony morphotypes, microcosms were vortexed for 1 minute and dilution plated on KB agar.

### Photography

Colonies were visualized under a Leica MS5 dissection microscope, and photographed with a VisiCam® 1.3 (VWR International). Microcosm photographs were taken with an iPhone 11. Photographs were cropped and, where noted in figure legends, the exposure and/or brightness uniformly altered in Preview (v11.0).

### Whole genome sequencing

Single colony isolates were grown to stationary phase in liquid KB (shaking) and genomic DNA isolated with a Qiagen DNeasy Blood and Tissue Kit (Qiagen). Extracted genomic DNA was checked for quality using agarose gel electrophoresis and a Nanodrop™ 3300 Fluorospectrometer. Whole genome re-sequencing was subsequently performed by the sequencing facility at the Max Planck Institute for Evolutionary Biology (Ploen, Germany), in three separate runs. The first run aimed to identify mutations in thirty SBW25-WS, PICF7-WS, and A506-Web isolates. 300 bp, paired-end reads were generated with a MiSeq Reagent Kit v3 (Illumina). Usable data was generated for 29 of 30 isolates; PICF7-WS5 was unsuccessful. PICF7-WS5 was re-sequenced in a second run, during which 150 bp, paired end reads were generated with a NextSeq 550 Output v2.5 Kit (Illumina). A third run was used to generate high coverage reads for A506 wild-type and one A506-Web isolate (A506-Web5). 150 bp, paired-end reads were generated with a NextSeq 550 Output v2.5 kit (Illumina). All sequencing reads are available at NCBI sequence read archive (Leinonen *et al.*, 2011), under BioProject number XXXXX-YYYYYY.

### Analysis of whole genome sequencing data for PICF7-WS and SBW25-WS

A minimum of 1 million reads *per* isolate were aligned to the PICF7 (GenBank CP005975.1; Martínez-García *et al.*, 2015) or SBW25 (RefSeq NC_012660.1; Silby *et al.*, 2009) reference genome sequences, using *breseq* (Deatherage and Barrick, 2014). A minimum mean coverage of 26.3 and 28.9 was obtained for the PICF7-WS and SBW25-WS genotypes, respectively. A list of possible mutations was generated for each genome (Table S3), and unique mutations were confirmed in the genome of interest by PCR amplification and Sanger sequencing. In isolates where no unique candidate mutations were initially identified by *breseq* (SBW25-WS6 and SBW25-WS8), WS mutations were identified by PCR amplification and Sanger sequencing of commonly mutated loci in WS types. For a detailed list of the WS-causing mutations, see Text S1.

### Analysis of whole genome sequencing data for A506

The first attempt at re-sequencing A506-Web isolates resulted in the prediction of >200 mutations *per* isolate by *breseq*, many of which were present in all ten isolates (Table S3). To clarify which, if any, of these predicted mutations were real and relevant for the Web phenotype, we sequenced our copy of A506 wild type and one Web isolate (A506-Web5) at high coverage. For each of these two samples, slightly under 6 million 150 bp, paired-end sequencing reads were obtained, and aligned to the NCBI reference sequences for the A506 chromosome (NC_017911.1; Loper *et al.*, 2012) and plasmid (NC_021361.1; Stockwell *et al.*, 2013) using *breseq* (Deatherage and Barrick, 2014). Mean coverages of 144.9 and 139.9 (A506 wild type chromosome and plasmid, respectively), and 151.1 and 111.9 (A506-Web5 chromosome and plasmid, respectively) were obtained, and ~100 mutations were predicted in each sample. All differences predicted in A506-Web5 were also predicted in A506 wild type, and no convincing additional differences were found in the lower coverage genome re-sequencing in any of the other Web isolates (Table S3).

### Transposon mutagenesis

Four *P. fluorescens* PICF7 WS genotypes (WS3, WS4, WS7, WS8; carrying mutations in *wspF*, *awsX*, *mwsR*, *wspF*, respectively) were subjected to a suppressor analysis *via* random mutagenesis with the IS-Ω-Km-/hah transposon (Jacobs *et al.*, 2003). The protocol previously described for the mutagenesis of *P. fluorescens* SBW25 (Giddens *et al.*, 2007) was used, with the exception that PICF7-WS recipient genotypes were not subjected to a heat shock prior to biparental conjugation. Approximately 15,600 transformant colonies from eleven independent conjugations were screened for loss of both the WS colony and 24-h mat forming phenotype. The transposon insertion site was determined in 100 suppressor mutants (Table S4). In selected transformants, the bulk of the transposon was deleted (“CreTn” types), leaving 189 bp at the insertion site and eliminating polar effects (Giddens *et al.*, 2007).

### Colony morphotype stability test

Single colonies of six strains (A506-Web1, A506-Web2, A506-Web3, A506-*wspF*-LoF, SBW25-WS4, and PICF7-WS2) were grown from glycerol stocks on KB agar. For each strain, three single colonies displaying the Web or WS phenotype were grown overnight in liquid KB. Overnight cultures were vortexed, dilution plated on KB agar. The number and morphotype of colonies from each culture was recorded. For each culture, between 45 and 300 colonies were observed, across three KB plates. The proportion of colonies showing the original morphotype (Web or WS) was recorded, and the mean and standard error calculated (Figure 4B; Raw Data File 1). In the case of strains carrying a *wspF* mutation (A506-*wspF*-LoF, SBW25-WS4, PICF7-WS2), no variation in colony morphology was observed. Parametric, one-tailed, one-sample *t*-tests were used to detect loss of Web colony morphology in A506-Web types. Analyses were performed in R (v3.6.0). Significance levels *0.05 *p*<0.01, **0.01<*p*<0.001.

### Genetic engineering in A506

Scar-free deletions were constructed in A506 using a similar protocol to that outlined for SBW25 (Zhang and Rainey, 2007). Briefly, deletion fragments were constructed by SOE-PCR (Ho *et al.*, 1989), ligated into the suicide vector pUIC3 (Rainey, 1999), and delivered into the relevant A506 background using a recombination-based, two-step allelic exchange protocol. The differences to the SBW25 protocol are (i) A506 was not subjected to a heat shock prior to the first recombination event, (ii) a higher concentration of Tc was used (50 μg ml^−1^) to select first recombinants, and (iii) cycloserine enrichment was not used to enrich for second recombinants; first recombinant types were simply grown in flasks and plated onto LB+Xgal agar to identify colonies that had lost the pUIC3 vector (*i.e.* white, Tc sensitive colonies). Presence of the deletion was confirmed using PCR and Sanger sequencing with primers outside the manipulation region. In each case, the final result is a strain in which the target sequence is cleanly deleted, leaving no trace of the pUIC3 vector. See Text S3 for more detail on each genotype, and the engineering process.

## ACKNOWLEDGEMENTS

We thank Paul Rainey, Jesús Mercado-Blanco, and Steven Lindow for the kind gifts of *P. fluorescens* SBW25, *P. simiae* PICF7, and *P. fluorescens* A506, respectively. We thank Frederic Bertels for assistance with building the *Pseudomonas* phylogenetic tree, and Gunda Dechow-Seligmann for technical assistance. Funds were received from the Max Planck Society, and the International Max Planck Research School for Evolutionary Biology (IMPRS-EB).

## AUTHOR CONTRIBUTIONS

AM and JG conceived the research, performed the experiments and data analyses, and wrote the manuscript.

## SUPPORTING INFORMATION

**Figure S1.**
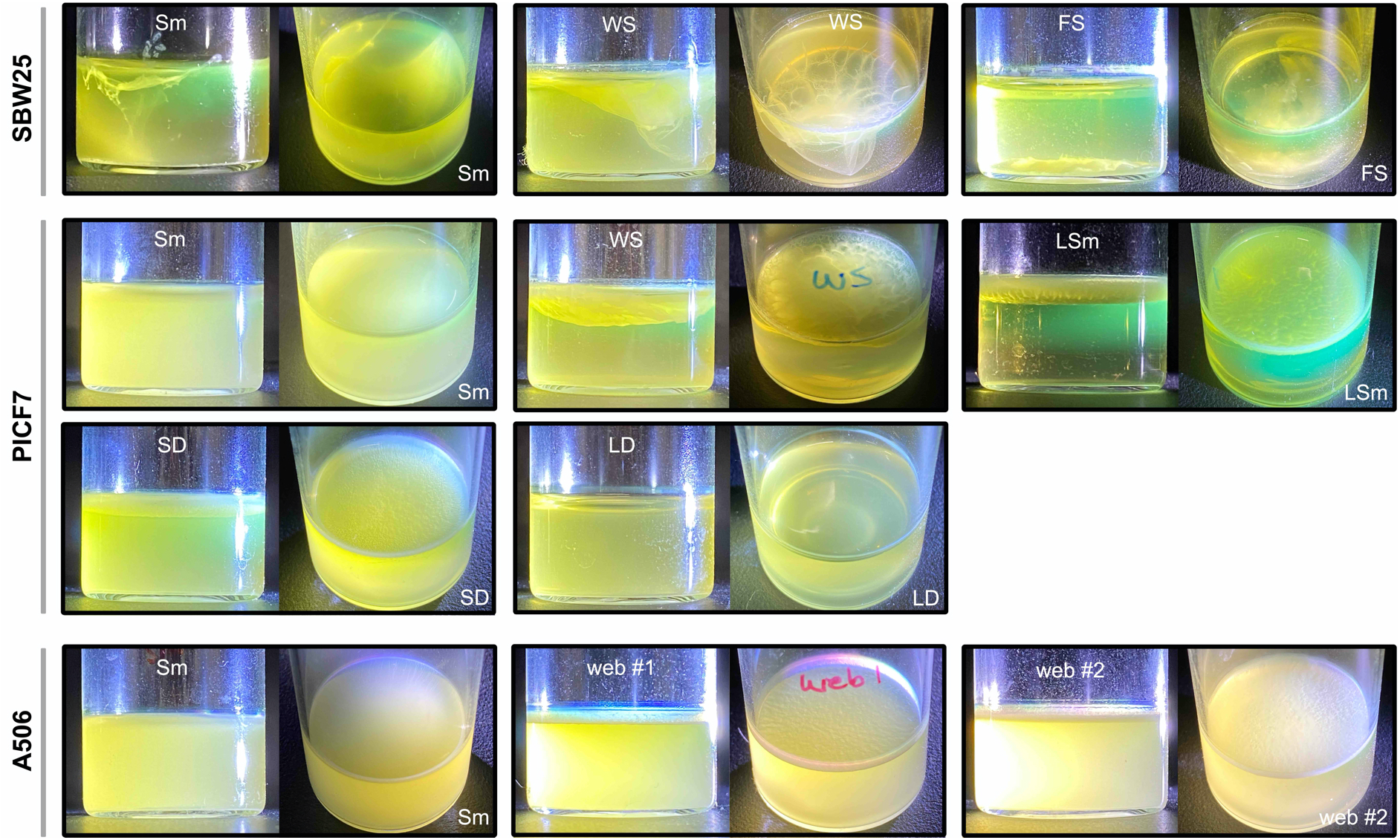
Five of the ten morphotypes isolated from 72-h microcosms have the emergent ability to form mats at the air-liquid interface within 24-h. Ten classes of colony phenotypes were observed arising from 72-h microcosms of SBW25, PICF7, and A506 (see Figure 2B). Of these, five classes (SBW25-WS, PICF7-WS, PICF7-LSm, PICF7-SD, and A506-Web) have the emergent ability to form mats within 24-h. The remaining five classes (SBW25-Sm, SBW25-FS, PICF7-Sm, PICF7-LD, and A506-Sm) do not form stable mats within 24 hours. All photographs taken at the same time point (24 h). Brightness of some images altered in Preview.

**Table S1. Strains, plasmids, transposons, and oligonucleotides used in this study.**

**Table S2. A comparison of regulatory and structural gene homologues in in *P. fluorescens* SBW25, *P. simiae* PICF7, and *P. fluorescens* A506.**

**Table S3. Mutations predicted by whole genome re-sequencing of ten SBW25-WS, ten PICF7-WS, and ten A506-Web isolates.** Illumina whole genome re-sequencing was performed for each isolate. This file contains (i) a summary of the raw read numbers and coverage information (“summary” tab), and (ii) a list of mutations predicted by *breseq* (Deatherage and Barrick, 2014), for each isolate (separated into “sbw25”, “picf7”, and “a506” tabs). WS mutations identified in PICF7-WS and SBW25-WS are further described in Text S1, and the possible duplications in A506 isolates (including wild type) are further described in Text S2.

**Table S4. Suppressor analysis of the PICF7-WS phenotype.** Genotype and phenotype details of 100 non-wrinkly transposon mutants from four PICF7-WS backgrounds (PICF7-WS3, WS4, WS7, and WS8; carrying mutations in *wspF*, *awsX*, *mwsR*, and *wspF*, respectively). Mutants were obtained by screening approximately ~15,600 transposon mutants from eleven independent conjugations for loss of the WS phenotype conferred by the WS mutation; each mutant forms smooth colonies, and cannot form a static microcosm mat within 24 hours. Mutants are classified according to the cellular function likely to be affected by the insertion. The precise genomic location of the transposon is indicated as the first base on the genomic forward strand downstream of the 3′ transposon terminus. For mutants of particular interest, a Cre-deletion (removing most of the transposon and thus eliminating polar effects) was obtained and analysed. Green highlighting shows transposon insertions in the *pel* biosynthetic locus; grey highlighting shows transposon insertions in loci involved in the regulation of Pel production (*wsp, aws*, or *mwsR*, depending on the WS mutation carried by the genotype being mutagenized).

**Text S1. Details of WS mutations in PICF7-WS and SBW25-WS.**

**Text S2. Analysis of the high coverage regions in *P. fluorescens* A506 isolates.**

**Text S3. Extended methods for the construction of scar-free mutations in *P. fluorescens* A506.**

## Notes

### Competing Interest Statement

The authors have declared no competing interest.

